# Genes encoding recognition of the *Cladosporium fulvum* effector protein Ecp5 are encoded at several loci in the tomato genome

**DOI:** 10.1101/759761

**Authors:** Michail Iakovidis, Eleni Soumpourou, Elisabeth Anderson, Graham Etherington, Scott Yourstone, Colwyn Thomas

**Author notes:** Private address, Greece (Michail Iakovidis). The Sainsbury Laboratory, University of Cambridge, Cambridge, CB2 1LR, UK. Private address, UK. Earlham Institute, Norwich Research Park, Norwich, NR4 7UZ, UK. Q^2^ Solutions, Morrisville, NC 27560, USA. Main co-corresponding author Dr Michail Iakovidis, private address, Greece. Co-corresponding author Dr Colwyn Thomas, Senior Lecturer, School of Biological Sciences, University of East Anglia, Norwich, Norfolk, NR4 7TJ, UK.

## Abstract

The molecular interactions between tomato and *Cladosporium fulvum* have been an important model for molecular plant pathology. Complex genetic loci on tomato chromosomes 1 and 6 harbor genes for resistance to *Cladosporium fulvum*, encoding receptor like-proteins that perceive distinct *Cladosporium fulvum* effectors and trigger plant defenses. Here, we report classical mapping strategies for loci in tomato accessions that respond to *Cladosporium fulvum* effector Ecp5, which is very sequence-monomorphic. We screened 139 wild tomato accessions for an Ecp5-induced hypersensitive response, and in five accessions, the Ecp5-induced hypersensitive response segregated as a monogenic trait, mapping to distinct loci in the tomato genome. We identified at least three loci on chromosomes 1, 7 and 12 that harbor distinct *Cf-Ecp5* genes in four different accessions. Our mapping showed that the *Cf-Ecp5* in *Solanum pimpinellifolium* G1.1161 is located at the *Milky Way* locus. The *Cf-Ecp5* in *Solanum pimpinellifolium* LA0722 was mapped to the bottom arm of chromosome 7, while the *Cf-Ecp5* genes in *Solanum lycopersicum* Ontario 7522 and *Solanum pimpinellifolium* LA2852 were mapped to the same locus on the top arm of chromosome 12. Bi-parental crosses between accessions carrying distinct Cf-Ecp5 genes revealed putative genetically unlinked suppressors of the Ecp5-induced hypersensitive response. Our mapping also showed that *Cf-11* is located on chromosome 11, close to the *Cf-3* locus. The Ecp5-induced hypersensitive response is widely distributed within tomato species and is variable in strength. This novel example of convergent evolution could be used for choosing different functional *Cf-Ecp5* genes according to individual plant breeding needs.

## INTRODUCTION

Plant-microbe interactions, characterized genetically by the gene-for-gene model (FLOR 1951), have typically been defined by the presence or absence of a pathogen avirulence gene and it’s cognate disease resistance (*R*) gene in the host which together determine the outcome of the interaction. The gene-for-gene model has been portrayed at the molecular level as being part of a larger dynamic evolutionary process known as the “zig-zag” model (JONES AND DANGL 2006), where *R* genes and pathogen effector/avirulence genes co-evolve in perpetual ‘boom-and-bust’ cycles (PRIESTLEY 1978). To date, effector-specific *R* genes have been predominantly confined to a single locus. Exceptions to this have been previously reported through convergent evolution of *R* genes in different species (Arabidopsis *RPM1* and soybean *RPG1*; Arabidopsis *RPS5* and wheat/barley *PBR1*) (ASHFIELD *et al.* 2014; CARTER *et al.* 2018), within the same species, but linked (Arabidopsis *RRS1*/*RPS4* and *RRS1B*/*RPS4B*) (SAUCET *et al.* 2015), and even within related species at unlinked loci (potato *Rpi-mcq1* and *Rpi-blb3*) (AGUILERA-GALVEZ *et al.* 2018).

The pathosystem of tomato (*Solanum lycopersicum*) and the leaf mould pathogen *Cladosporium fulvum* is a well-studied model of gene-for-gene interactions and plant disease resistance gene evolution (RIVAS AND THOMAS 2005; DE WIT 2016). The fungus is considered an asexual non-obligate biotroph of the *Mycosphaerellaceae* family that infects plants via conidia that settle on the abaxial leaf side, germinating and entering the plant through open stomata, leading to reduced respiration, defoliation and even host death (THOMMA *et al.* 2005). The fungus abundantly secretes effector proteins in the leaf apoplast and several of these effectors can be recognized by cognate *R* genes in specific tomato accessions. The tomato *R* genes (designated *Cf* genes, for resistance genes to *Cladosporium fulvum*) encode receptor-like proteins (RLPs) that localize to the plasma membrane and contain *e*xtracellular *l*eucine-*r*ich *r*epeats (eLRRs), a membrane spanning domain, and a non-signaling short cytoplasmic domain. This system has acted as a model for investigating the structure and evolution of plant disease resistance gene loci (THOMAS *et al.* 1998; RIVAS AND THOMAS 2005; DE WIT *et al.* 2012; LIN *et al.* 2014; DE WIT 2016). To date, these genes have been mapped to just two chromosomal segments in the tomato genome. The *Milky Way* (*MW*) locus on the short arm of chromosome 1, and several genetically linked loci (like *ORION; OR*) containing functional *Cf* genes, encode a large number of genes with distinct recognition specificities including *Cf-4*, *Hcr9-4E*, *Cf-9*, *Hcr9-9B*, *Cf-19*, *Cf-Ecp1*, *Cf-Ecp2*, *Cf-Ecp3*, *Cf-Ecp4, Cf-Ecp5* (JONES *et al.* 1994; THOMAS *et al.* 1997; PARNISKE *et al.* 1999; TAKKEN *et al.* 1999; HAANSTRA *et al.* 2000; PANTER *et al.* 2002; KRUIJT *et al.* 2004; SOUMPOUROU *et al.* 2007; ZHAO *et al.* 2016). These genes were designated as *Hcr9s* (*h*omologs of *Cladosporium r*esistance gene 9). Another complex locus (*Cf-2*/*Cf-5*) encoding *Cf* genes has been characterized on the short arm of chromosome 6, that includes *Cf-2.1, Cf-2.2* and *Cf-5* (DIXON *et al.* 1996) with their genes having been designated *Hcr2s* (*h*omologs of *Cladosporium r*esistance gene 2) respectively (RIVAS AND THOMAS 2005). However, the chromosomal locations of several other *Cf* genes, which may comprise new complex loci, have not yet been reported (RIVAS AND THOMAS 2005; DE WIT 2016).

Here, we deployed genetic mapping to investigate the genetics of the tomato hypersensitive response-type (HR) or cell death to *C. fulvum* extracellular protein 5 (Ecp5). Ecp5 is 115 aa long with 6 cysteine residues and is also one of the least polymorphic *C. fulvum* effectors (HAANSTRA *et al.* 2000; STERGIOPOULOS *et al.* 2007; DE WIT *et al.* 2009). Similarly to *C. fulvum* avirulence proteins (Avrs), Ecps are secreted by all *C. fulvum* strains and are known to play an essential role in infection, since their deletion results in decreased virulence of *C. fulvum* on tomato, to which Ecp5 is no exception (DE WIT *et al.* 2009; DE WIT *et al.* 2012; MESARICH *et al.* 2017). A closely related fungus to *C. fulvum*, *Dothideomycetes septosporum* contains only a pseudogenised Ecp5 homologue, despite having functional homologues of other *C. fulvum* effectors like Avr4 and Ecp2 (DE WIT *et al.* 2012).

A single gene controlling the Ecp5-induced HR in tomato was previously designated *Cf-Ecp5* in the line *S. lycopersicum* G1.1161 (introgressed from *S. pimpinellifolium*) and mapped at the *AURORA* locus (*AU*), proximal to *MW* (HAANSTRA *et al.* 2000). Additional *S. pimpinellifolium* and *S. lycopersicum* accessions appeared to also carry a single dominant *Cf-Ecp5* gene and develop a HR following inoculation with recombinant potato virus X (*PVX*) strains expressing Ecp5 (HAANSTRA *et al.* 1999; HAANSTRA *et al.* 2000). It was presumed that all of them carried a similar *Cf-Ecp5* gene at *AU* and a transposon tagging strategy was deployed to isolate it from *S. lycopersicum* Ontario 7522 (Ont7522). However, unexpected genetic ratios during initial crossing lead to further investigation and reassignment of *Cf-Ecp5* in G1.1161 to the *MW* locus. Three other *Cf-Ecp5* loci from *S. pimpinellifolium* LA0722 and LA2852, and *S. lycopersicum* Ont7522 were shown to be genetically unlinked to *MW*. We used AFLP bulked segregant analysis and mapped *Cf-Ecp5* in LA0722 to the bottom arm of chromosome 7 and *Cf-Ecp5* in LA2852 and Ont7522 to the top arm of chromosome 12. Differential cell death symptoms were also observed amongst the *S. pimpinellifolium* and *S. lycopersicum Cf-Ecp5*-carrying accessions and allelism crosses between them revealed distinct Ecp5-dependent host regulators. We screened a total of 139 domesticated and wild tomato accessions for Ecp4- and Ecp5-induced HR and identified multiple accessions carrying *-Cf-Ecp4* or *Cf-Ecp5* genes that could be used in future studies. We found that *Cf-Ecp5* was distributed in a wider geographical region than *Cf-Ecp4*. Lastly, we report the chromosomal locations of *Cf-11*, which maps to the top arm of chromosome 11, close to the previously reported Cf-3 gene (KANWAR *et al.* 1980). All in all, this study revealed a unique example of multiple convergently evolved tomato loci in different accessions associated with the HR to *C. fulvum* Ecp5 effector that have not been previously reported in plant-pathogen interactions.

## MATERIALS AND METHODS

### Plant and fungal materials

Seeds from core collections of wild and domesticated tomato species were obtained from the C.M. Rick *Tomato Genetics Resource Center*, University of California, Davis, USA. These species included *Solanum arcanum*, *Solanum cheesmaniae*, *Solanum chilense*, *Solanum chmielewskii*, *Solanum corneliomulleri*, *Solanum galapagense*, *Solanum habrochaites*, *Solanum huaylasense*, *Solanum lycopersicoides*, *Solanum lycopersicum*, *Solanum neorickii*, *Solanum ochranthum*, *Solanum pennellii*, *Solanum peruvianum* and *Solanum sitiens*. Additional stocks containing previously characterised *Cf-Ecp* genes were obtained from Dr M. H. A. J. Joosten and Dr P. Lindhout (University of Wageningen) including the accessions *S. pimpinellifolium* LA1683 (*Cf-Ecp4*) and *S. lycopersicum* G1.1161 containing the introgressed *S. pimpinellifolium* gene *Cf-Ecp5* (HAANSTRA *et al.* 2000).

A number of other stocks were used in this study for the genetic mapping of *Cf-Ecp5* genes. The following lines were supplied by The Sainsbury Laboratory, Norwich, UK; *S. lycopersicum* variety ‘*Moneymaker*’ (MM), which contains no *Cf* genes (Cf0); the FT33 line (ROMMENS *et al.* 1992), which contains a T-DNA located 3 centimorgans (cM) proximal to the *MW* locus on the short arm of chromosome 1 and which harbors the maize transposon *Dissociation* (*Ds*) that carries the *Escherichia coli uidA* gene (*GUS*); line M18 contains an insertion of *Ds* in the *Cf-9* gene (*Ds*∷*Cf-9*) at the *MW* locus (Jones *et al.*, 1994). Forty-nine *S. pennellii* introgression lines (ILs) in the *S. lycopersicum* M82 background (ESHED AND ZAMIR 1994) were used for AFLP mapping of *Cf-Ecp5* genes to defined chromosomal intervals as described previously (THOMAS *et al.* 1995). *C. fulvum* race 3 (CAN 84) and race 2.4.5.9.11 were obtained from Dr MHA Joosten at the University of Wageningen, Netherlands. *C. fulvum* race 4 GUS (OLIVER *et al.* 1993) and race 5 were maintained and prepared for plant infections as described previously (HAMMOND-KOSACK *et al.* 1994). The *S. lycopersicum* stock Ont7716 and near-isogenic lines (NILs) containing single introgressed *Cf* genes were obtained from The Sainsbury Laboratory in Norwich, UK (TIGCHELAAR 1984).

### DNA preparation and marker sequence analysis

DNA was prepared from individual tomato lines and bulked segregant pools as described previously (THOMAS *et al.* 1995; SOUMPOUROU *et al.* 2007). AFLP analysis of tomato genomic DNA was performed as described previously (THOMAS *et al.* 1995). Cleaved amplified polymorphic sequence (CAPS) and simple sequence repeat (SSR) analyses were all performed as described previously (SOUMPOUROU *et al.* 2007).

### DNA gel blot analysis

Two to five μg of tomato DNA was digested at 37ºC for 16 h, extracted with phenol-chloroform and ethanol precipitated. DNA was electrophoresed for 20 h at 2.5 V/cm in 0.8% w/v agarose gels in a vertical gel apparatus using 40 mM Tris-acetate, pH 7.9, 1 mM EDTA as running buffer. Nucleic acids were transferred to Hybond-N membrane (Amersham) and cross-linked by irradiation with UV light. Filters were hybridised with ^32^P-labelled probes at 65ºC for 16-20 h and washed four times for 15 min in 2x SSC (1x SSC is 0.15 M sodium chloride, 0.015 M sodium citrate) and 1% w/v SDS at 65ºC, and for 30 min in 0.2x SSC and 0.1% w/v SDS at 65ºC.

### Assaying *Cf-Ecp5* function by infiltration with recombinant potato virus X

*Cf-Ecp5* function can be assayed in several ways based on expression of the *C. fulvum* Ecp5 protein in plants. In a genetic test, the F_1_ progeny of crosses between *Cf-Ecp5* containing lines and MM plants stably expressing Ecp5 exhibit a characteristic seedling lethal phenotype. Transgenic MM lines expressing the *C. fulvum* Ecp5 protein (MM-Ecp5) were described previously (SOUMPOUROU *et al.* 2007). Alternatively, *Cf-Ecp5* function can be determined by delivering ECP5 in the form of recombinant *Potato Virus X*, expressing ECP5 (SOUMPOUROU *et al.* 2007). Plants expressing *Cf-Ecp5* exhibit systemic necrosis following virus replication and spread. Stocks of *A. tumefaciens* GV3101 were streaked onto Luria-Broth (LB) agar plates containing 40 μg/ml kanamycin. Colonies were selected and cultured in 10 ml LB medium containing 40 μg/ml kanamycin on a shaking platform at 28⁰C overnight. The cultures were centrifuged at 2000 x *g* and the pellets re-suspended in a solution containing 1x Murashige & Skoog salts and 2% w/v sucrose. To initiate the expression of *vir* genes, acetosyringone (3’-5’-Dimethoxy-4’-hydroxy–acetophenone) was added to a final concentration of 150 μM and the bacteria were left at room temperature for 3 hr. Bacteria were infiltrated into a single cotyledon of 7-10 day old seedlings. Plants containing the cognate *Cf* gene showed signs of systemic necrosis at 7-8 days post-infection (dpi) (THOMAS *et al.* 1997; SOUMPOUROU *et al.* 2007).

### Marker analysis

AFLP analysis of bulked-segregant pools for identifying markers linked *in trans* with various *Cf-Ecp5* genes was performed as described previously (THOMAS *et al.* 1995). Generation of CAPS (Cleaved Amplified Polymorphic Sequence) markers between specific haplotypes involved PCR amplification of sequences based on the tomato EXPEN2000 genetic map (http://solgenomics.net/) and sequencing of haplotype-specific products to detect sequence polymorphisms. PCR was carried out in a total volume of 20 μmu;l containing 10 mM Tris-HCl, pH 8.0, 1.5 mM MgCl_2_, 0.2 mM dNTPs and 0.2 units Taq DNA polymerase, 50 ng genomic DNA and 1 μM of each primer. PCR products were purified using Qiaspin (Qiagen) columns and were sequenced on a 3730XL sequencer. Details of primers and CAPS markers used in this study are shown in Table S1. In reference to previous studies, genetic distances were estimated by converting recombination fractions using the Haldane function.

### Bioinformatic analysis to detect LRR-encoding genes in defined regions of the tomato genome

To identify candidate RLP genes for *Cf-Ecp5* on tomato chromosomes 7 and 12 target regions delimited by flanking Conserved Ortholog Set II (COSII) markers were defined. For each target region protein sequences for the two markers were extracted from the TAIR website (http://www.arabidopsis.org/) and a TBLASTN search was carried out against version 2.30 of the ‘Heinz 1706’ tomato genome (CONSORTIUM 2012). Using these results we defined target regions on chromosomes 7 and 12 which are shown in Table S7. The nucleotide sequence for each target region was extracted and six-frame translations were generated using the EMBOSS tool TRANSEQ (RICE *et al.* 2000). The *β*-sheets of Cf proteins (and other RLP and eLRR-RK proteins) contain highly conserved leucine-rich repeat motifs (LxxLxLxxNxLxGxIP) (RIVAS AND THOMAS 2005). Using the six-frame translations, a Perl regular expression search was carried out to detect motifs with this canonical structure. From this search we found 13 motifs within the chromosome 7 target and 35 within the chromosome 12 target. Of the 48 sequences, 34 were unique. The unique sequences were then taken and using MEME a mixture model was created (BAILEY AND ELKAN 1994) to form a search model for LRR motifs. Before using the model to search for unknown RLPs within the two target regions, the model was tested on a region of the tomato genome known to encode RLPs. The tomato resistance genes *Cf-2* and *Cf-5* are located on chromosome 6 (JONES *et al.* 1993) and the nucleotide sequence for *Cf-5* (AF053993) was used in a BLAST analysis of the tomato genome. Two highly homologous genes were found at the predicted location of the ‘Heinz 1706’ sequence between positions 2138887 – 2142639 and 2161278 – 2167796. An 18 kb gap separates these two genes, so the target area included the gene-spanning region plus an additional 18 kb on either side was used to test our model. Using FIMO (part of the MEME Suite) (BAILEY *et al.* 2009), we then searched for LRR-like motifs in this region with a FIMO default p-value of 1e^-4^. Sixty-four LRR-like motifs were identified in the target area, fifty-six within the two *Cf-2/Cf-5* homologs, and eight outside it. The lowest p-value of the presumed eight false-positives outside of the *Cf-2/Cf-5* homologs was 1.06e^-05^. When all 64 LRR-like motifs were filtered on this p-value, we were left with 49 LRR-like motifs within the two homologs, and none outside. We then used FIMO with a q-value threshold of 1.06e^-05^ to identify LRR-encoding motifs within the chromosome 7 and 12 target regions and regions with consecutive blocks of LRR-encoding motifs as candidate LRR-encoding genes.

### Data availability

All tomato seed stocks, bacterial and fungal strains are available upon request. The GSA Figshare portal was used to upload supplemental data files.

## RESULTS

### The *Cf-Ecp5* gene from *S. pimpinellifolium* G1.1161 maps at *MW*

We previously reported the mapping of two other *S. pimpinellifolium* genes (*Cf-Ecp1* and *Cf-Ecp4*) at *MW*, where the majority of *Cf* genes have been mapped (SOUMPOUROU *et al.* 2007). We also identified recombinants between the FT33 locus (which contains a T-DNA insertion harboring *Ds:GUS* and located 3 cM proximal to *MW*) and four *Cf-Ecp* genes we targeted for cloning. The observed recombination frequencies between the FT33 locus and genes located at *MW* (*Cf-Ecp1* and *Cf-Ecp4*), and the *Orion* (*OR*) locus (*Cf-Ecp2* being 10 cM proximal to *MW*), appeared consistent with their reported map locations (HAANSTRA *et al.* 1999; SOUMPOUROU *et al.* 2007). The *Cf-Ecp5* gene in *S. pimpinellifolium* G1.1161 was originally mapped at the *Aurora* (*AU*) locus, 3 cM proximal to *MW*, and therefore close to the FT33 locus (HAANSTRA *et al.* 2000). However, in our study the observed recombination frequency between FT33 and *Cf-Ecp5* was higher than expected (3.67%) (SOUMPOUROU *et al.* 2007), and similar to the recombination frequencies observed between FT33 and *Cf* genes at *MW.* This result suggested that *Cf-Ecp5* is either located at *MW* or proximal to FT33.

An allelism test was performed where G1.1161 was crossed with line M18 which contains a stable *Ds:GUS*-tagged allele of *Cf-9* (JONES *et al.* 1994). The F_1_ plants were then test-crossed to Cf0:*Ecp5*, Cf0, or *S. pennellii* LA0716 to generate segregating populations. The resulting progeny were screened for GUS activity and either the seedling lethal phenotype (crosses to Cf0:*Ecp5*) or a systemic HR following agro-infiltration with *PVX*:*Ecp5* (crosses to Cf0 or LA0716). Out of 734 progeny, no recombinants were observed between *Cf-Ecp5* and the *Ds*∷GUS-tagged *Cf-9*, confirming they are either allelic or very tightly linked (Table 1). This result is consistent with *Cf-Ecp5* being encoded by an *Hcr9* at *MW* in *S. pimpinellifolium* G1.1161 and was therefore designated *Cf-Ecp5.1*.

**Table 1.** The Cf-Ecp5 gene in S. *pimpinellifolium* G1.1161 is allelic or very tightly linked to the *Cf-9* gene at MW. Test cross progeny were screened using a standard histological procedure for the presence or absence of GUS activity (GUS or no GUS) and for the presence or absence of Cf-Ecp5 (<u>N</u>ecrotic or <u>w</u>ild <u>t</u>ype respectively) based on the development of either a systemic necrosis phenotype following agro-infiltration with *PVX:Ecp5* (crosses to Cf0 and LA0716) or a seedling lethal phenotype (crosses to Cf0:Ecp5).

| **Phenotype** | **Cf0 x (G1.1161 x M18)** | **LA0716 x (G1.1161 x M18)** | **Cf0:*Ecp5* x (G1.1161 x M18)** |
| --- | --- | --- | --- |
| GUS:wt | 175 | 79 | 109 |
| no GUS:N | 201 | 62 | 108 |
| GUS:N | 0 | 0 | 0 |
| no GUS:wt | 0 | 0 | 0 |
| **Recombinant fraction** | **0%** | **0%** | **0%** |

### The *Cf-Ecp5* gene in *S. pimpinellifolium* LA2852 and *S. lycopersicum* Ont7522 map to chromosome 12

We assumed that *Cf-Ecp5* genes previously identified in other tomato accessions (LA0722, LA1689, LA2852, Ont7522) would map to the same chromosomal location as *Cf-Ecp5.1* at *MW* (LAUGÉ *et al.* 2000). Our preliminary analysis demonstrated that *Cf-Ecp5* assorted independently of the FT33 locus in these accessions. To map these genes we identified *S. pennellii* markers linked *in trans* to each gene which could then be located on *S. pennellii* introgression lines (ILs) (ESHED AND ZAMIR 1994). *S. pimpinellifolium* LA2852 and *S. lycopersicum* Ont7522 were each crossed to *S. pennellii* LA0716 and testcrossed to Cf0, to generate segregating populations. Testcross progeny were agro-infiltrated with *PVX*:*Ecp5* and scored for Ecp5-induced HR. For each cross, DNA bulked segregant pools were created from the cotyledons of forty testcross plants exhibiting either wild-type (wt) or Necrotic (N) symptoms (Figure 1A, 1B).

**FIGURE 1.**
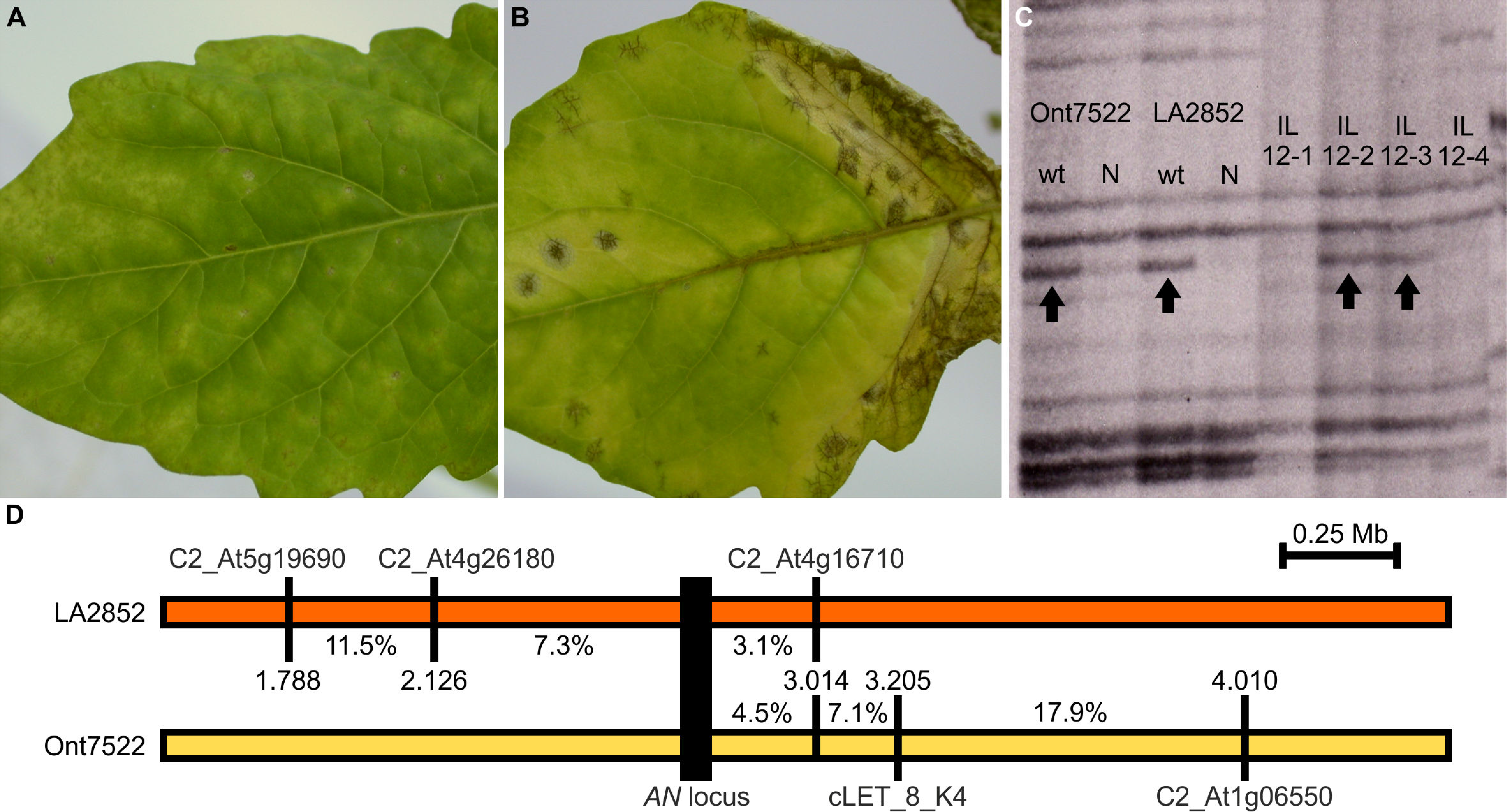
The *Cf-Ecp5* gene in *S. pimpinellifolium* LA2852 and *S. lycopersicum* Ont7522 map to the top arm of chromosome 12 in tomato. (A) Visible symptoms of wild type *PVX* and (B) systemic HR cell death in LA2852 accession triggered after *PVX*:*EV* and *PVX*:*Ecp5* agroinfiltration respectively. Pictures were taken at 14 dpi. (C) AFLP mapping of *S. pennellii* marker 91R31-M48 linked *in trans* with *Cf-Ecp5* in Ont7522 and LA2852 in the chromosomal interval between IL12-2 and IL12-3. Each phenotypic bulk is shown under the “wt” and “N” labels for each accession (“wt” = wild type, “N” = Necrotic). *S. pennellii* ILs (12-1 to 12-4) covering the entire chromosome 12 are shown. (D) Genetic map of the chromosome 12 region at the *Cf-Ecp5* locus in LA2852 (orange box) and Ont7522 (yellow box). The interval of *Cf-Ecp5* in LA2852 was delimited by genotyping 48 wild-type Cf0 x LA2852 F_2_ plants with three markers: C2_At5g19690 (11/96 recombinant gametes), C2_At4g26180 (7/96 recombinant gametes), and C2_At4g16710 (3/96 recombinant gametes). The *Cf-Ecp5* in Ont7522 was delimited by genotyping of 56 wild-type Cf0 x Ont7522 F_2_ plants with also three markers: At1g06550 (20/112 recombinant gametes), cLET_8_K4 (8/112 recombinant gametes), C2_At4g16710 (5/112 recombinant gametes). Distances are shown in megabases (Mb) and the location of each marker on the chromosome is also shown as Mb coordinates. The recombination fraction (%) between the *AN* locus (black box) and each marker is shown as well for both haplotypes.

The *S. pennellii* markers linked *in trans* were similar for LA2852 and Ont7522 phenotypic bulks, suggesting the two *Cf-Ecp5* genes are located in the same chromosomal region. Three AFLP markers were selected to be characterised for the Ont7522 population using 12 bulks of *S. pennellii* ILs (each representing an entire tomato chromosome), with all three markers mapping to chromosome 12 (Figure S1). The chromosomal interval for *Cf-Ecp5* in both LA2852 and Ont7522 was then localized at the overlap between IL12-2 and IL 12-3 (Figure 1C).

To further delimit the chromosomal location of these genes, two sets of 200 F_2_ plants from a Cf0 x LA2852 and a Cf0 x Ont7522 cross were agro-infiltrated with *PVX*:*Ecp5* and scored for wild-type symptoms at 14 dpi. Both sets of data were consistent with a 3:1 segregation ratio for a dominant monogenic trait (Cf0 x LA2852 F_2_, 152:48 N:wt, *p*-value (χ^2^) = 0.816; Cf0 x Ont7522 144:56 N:wt, *p*-value (χ^2^)=0.497). CAPS markers were developed to distinguish the haplotypes used in these crosses (Table S1) primarily using COSII markers on the target chromosome as detailed at the Sol Genomics Network (http://solgenomics.net/). For genetic map construction, three informative markers were identified for LA2852 haplotype (C2_At5g19690, C2_At4g26180, C2_At4g16710) that flank the locus on both sides (Figure 1D), and for Ont7522 haplotype, the three informative markers that were identified, flanked the locus on one side only (At1g06550, cLET_8_K4, C2_At4g16710) (Figure 1D). Only one marker was useful in mapping both genes (C2_At4g16710), which is located proximal to both *Cf-Ecp5* genes at a similar distance (3.1% and 4.5% recombination respectively). This data suggest that the *Cf-Ecp5* genes in LA2852 and Ont7522 are located in a 888 kb region (2.126-3.014 Mb) and are either very closely linked or allelic, and therefore define a new *Cf* locus that we designated *ANDROMEDA* (*AN*), following nomenclature adopted in previous studies (PARNISKE *et al.* 1997; HAANSTRA *et al.* 2000). The genes in *S. pimpinellifolium* LA2852 and *S. lycopersicum* Ont7522 were then designated *Cf-Ecp5.12p* and *Cf-Ecp5.12l* respectively. The *AN* locus resides in a region that appears to contain many RLPs (KANG AND YEOM 2018).

### *Cf-Ecp5* in *S. pimpinellifolium* LA0722 maps to chromosome 7

To investigate if *Cf-Ecp5* in *S. pimpinellifolium* LA0722 is also located at *MW*, we used the same strategy described above for line G1.1161. Two hundred and ten progeny of a (M18 x LA0722) x Cf0 cross were assayed for GUS activity and then agro-infiltrated with *PVX*:*Ecp5* to determine their *Cf-Ecp5* genotype. The progeny segregated in a 1:1:1:1 ratio for all four possible phenotypic classes (51:47:60:52 GUS/N: GUS/wt: no GUS/wt: no GUS/N, *p*-value(χ^2^) = 0.843). This result indicates that *Cf-Ecp5* in LA0722 assorts independently of *MW*. A second allelism test involved a cross between a line containing *Cf-Ecp5.12p* and *S. pimpinellifolium* LA0722 that revealed putative genetic elements controlling HR (Table 2). To provide additional molecular evidence of independent assortment, 200 F_2_ plants from a Cf0 x LA0722 cross were agro-infiltrated with *PVX*:*Ecp5* and 59 wild-type individuals were identified, suggesting dominant monogenic inheritance (141:59 N:wt, *p*-value(χ^2^) = 0.312). We genotyped 59 wild-type plants with markers linked to known *Cf* gene loci on chromosomes 1, 11, and 12 (Table S2), and the data were indicative of an independently assorting *Cf-Ecp5* gene in LA0722 that defined a new locus, *CENTAURI* (*CE*).

**Table 2.** Unexpected suppression of Ecp5-related HR in crosses between different Cf-Ecp5-carrying accessions. Data from different populations following agroinfiltration with *PVX:Ecp5*. Phenotypes were recorded at 10-14 dpi. G1.1161 carries a *Cf-Ecp5* gene on chromosome 1, Ont7522 and LA2852 carry a *Cf-Ecp5* on chromosome 12, while Cf0 and LA0716 lack *Cf-Ecp5*. Population B851 lacked *Cf-Ecp5*.12p, while B852 and B853 carried it. In this study, LA2852 was usually crossed to Cf0 initially to form a bridged cross due to difficulties in direct crosses to other accessions. The cross, phenotypes (<u>w</u>ild-<u>t</u>ype or <u>N</u>ecrotic), expected ratios and p-values are shown (p-values were calculated in each cross based on the expected ratio if there was no suppression for dominant traits; ‘⭙↓’ = selfing; statistical significance was calculated based on chi-square tests: ‘***’=P≤.0.001; ‘**’=P≤.0.01; ‘*’=P≤.0.05).

| Cross | wild-type | Necrotic | Expected Ratio (wt:N) | *p*-value($\chi^2$) |
| --- | --- | --- | --- | --- |
| (Ont7522 x (Cf0 x LA2852)) ⊗↓ (B851) | 59 | 133 | 1:3 | 0.211 |
| (Ont7522 x (Cf0 x LA2852)) ⊗↓ (B852) | 42 | 362 | 0:1 | 0*** |
| (Ont7522 x (Cf0 x LA2852)) ⊗↓ (B853) | 41 | 364 | 0:1 | 0*** |
| (G1.1161 x (Cf0 x LA2852)) ⊗↓ | 121 | 196 | 1:15 | 0*** |
| (Ont7522 x LA0716) ⊗↓ | 74 | 81 | 1:3 | 3.159e-05*** |
| (Cf0 x LA2852) x LA0716 | 124 | 52 | 1:1 | 8.836e-05*** |

To locate the *CE* locus, we used the previously described bulked segregant AFLP mapping strategy for *Cf-Ecp5.12p* and *Cf-Ecp5.12l*. Progeny from a (LA0722 x M18) x LA0716 cross were testcrossed to Cf0 and families segregating for *Cf-Ecp5* were identified. The selected progeny were scored for Ecp5-induced HR and DNA bulks were created from 30 plants exhibiting either wild-type or necrotic symptoms. AFLP analysis was used to identify markers linked *in trans* with *Cf-Ecp5* at *CE* (Figure S2). Analysis of IL chromosome pools showed that the *Cf-Ecp5* in LA0722 maps on tomato chromosome 7 in the interval defined by IL7-4 (Figure 2A, 2B). As a consequence, this genetically distinct *Cf-Ecp5* gene in *S. pimpinellifolium* LA0722 was designated *Cf-Ecp5.7.* To construct a genetic map, 200 F_2_ plants from a Cf0 x LA0722 cross were agro-infiltrated with *PVX*:*Ecp5* and DNA was isolated from all individuals that exhibited wild-type symptoms at 14 dpi. These data are consistent with a 3:1 segregation ratio for dominant monogenic traits (Cf0 x LA0722 F_2_, 142:58 N:wt, *p*-value(χ^2^) = 0.368). We used four CAPS markers for genotyping the 58 wild-type F_2_ individuals: C2_At5g14520, C2_At1g78620, C2_At3g15290, and C2_At4g26750, which delimited *Cf-Ecp5.7* in a 6.1 Mb region (58.902-64.993 Mb) (Figure 2C). This region contains many different types of *n*ucleotide-binding domain *l*eucine-rich *r*epeat-containing genes (NLRs), but few RLPs, in the *Solanum lycopersicum* genome (JUPE *et al.* 2013; ANDOLFO *et al.* 2014; WEI *et al.* 2016; KANG AND YEOM 2018).

**FIGURE 2.**
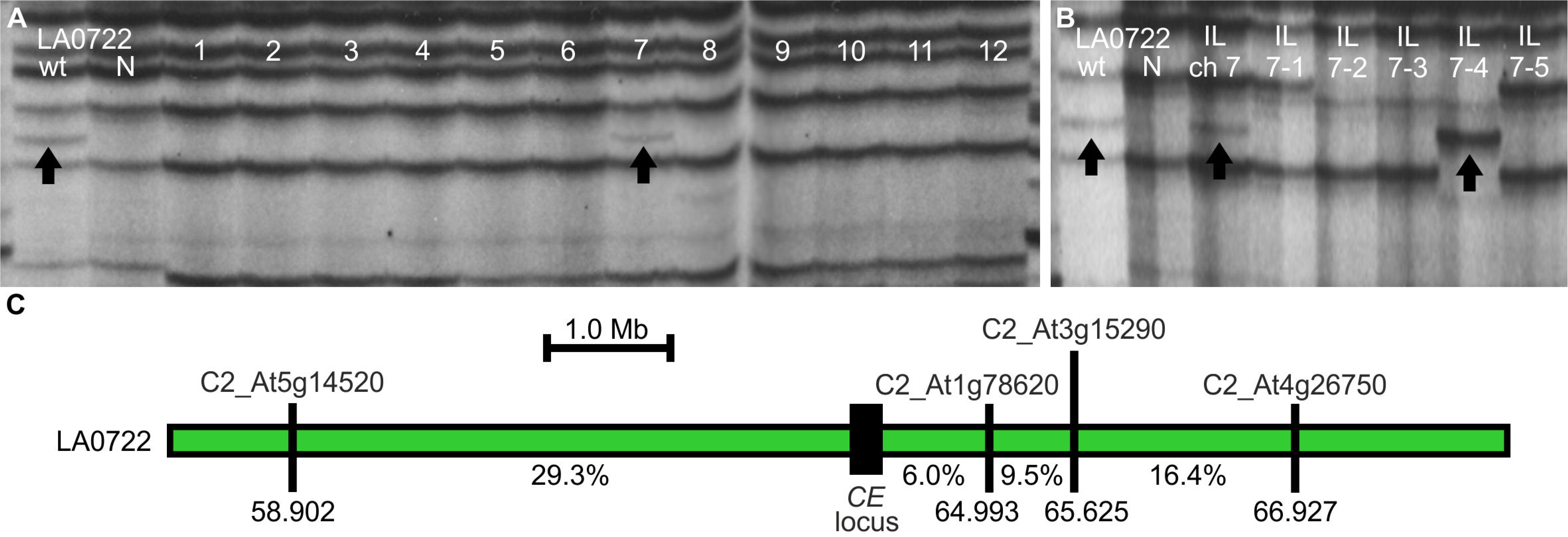
The *Cf-Ecp5* gene in *S. pimpinellifolium* LA0722 maps on the bottom arm of chromosome 7 in tomato. (A) Mapping of *S. pennellii* AFLP marker 91R30-M88 linked *in trans* with *Cf-Ecp5* in LA0722 alongside bulks of ILs that represent entire tomato chromosomes. Each phenotypic bulk is shown under the “wt” and “N” labels for each accession (“wt” = wild type, “N” = Necrotic). Tomato chromosome numbers (1-12) for IL bulks are shown for each lane. (B) AFLP mapping of *S. pennellii* marker 91R30-M88 linked *in trans* with *Cf-Ecp5* in LA0722 in the chromosomal interval IL7-4. *S. pennellii* bulk IL for chromosome 7 (IL ch 7) and individual ILs (7-1 to 7-5) covering the entire chromosome 7 are shown. (C) Genetic map of the chromosome 7 region at the *Cf-Ecp5* locus in LA0722 (green box). The interval was delimited by genotyping 58 wild-type Cf0 x LA0722 F_2_ plants with four markers: C2_At5g14520 (34/116 recombinant gametes), C2_At1g78620 (7/116 recombinant gametes), C2_At3g15290 (11/116 recombinant gametes), and C2_At4g26750 (19/116 recombinant gametes). Arrows indicate AFLP markers linked *in trans* to *Cf-Ecp5* in LA0722. Distances are shown in megabases (Mb) and the location of each marker on the chromosome is also shown as Mb coordinates. The recombination fraction (%) between the *CE* locus (black box) and each marker is shown.

### The Ecp4- and Ecp5-induced HR is conserved in the wild tomato germplasm

A previous genus-wide screen of wild tomato accessions for the presence of the well-characterised *Cf-4* and *Cf-9* genes demonstrated these two genes are conserved throughout the *Solanum* genus (KRUIJT *et al.* 2005). Previous studies have identified *Cf-Ecp5* genes in *S. pimpinellifolium* (G1.1161, LA0722, LA1689, LA2852) and *S. lycopersicum* (Ont7522) (LAUGÉ *et al.* 2000). In this study, we tested different accessions from multiple species that could trigger an Ecp4- or Ecp5-induced HR (Table S3). *PVX*-mediated delivery of *C. fulvum* Ecps provides a fast and reliable screen for the presence of their cognate *Cf* genes in wild *Solanum* species. Despite the low germination rate in many seed stocks, of the 139 accessions tested, *Cf-Ecp5* was detected in 16 accessions of 10 different tomato species (Figure 3). Five species appeared to lack *Cf-Ecp5* (*S. galapagense*, *S. habrochaites*, *S. ochranthum*, *S. lycopersicoides* and *S. sitiens*), but this may reflect the small sample size tested and low germination rates (Table S3). Most accessions containing *Cf-Ecp5*, included also non-responsive plants that exhibited wild-type *PVX* symptoms, indicating that they were segregating and heterozygosity might be common in these. Across all the Ecp5-responsive accessions, HR symptoms appeared variable in strength, similar to the five accessions investigated in this study (G1.1161, LA0722, LA1689, LA2852, Ont7522) (Figure S3). Despite all of them being inherited as dominant monogenic traits, differential observations in *Cf-Ecp5*-triggered HR symptoms in both agro-infiltrations with *PVX*:*Ecp5* and crosses to transgenic Cf0:*Ecp5* plants in G1.1161, LA0722, LA2852 and Ont7522 were indicative of the distinctiveness of their *Cf-Ecp5* genes (Figure S3). The Ecp5-induced HR in LA2852 and Ont7522 were the strongest, followed by LA0722 and LA1689, and lastly G1.1161, which exhibited the weakest HR symptoms (Figure S3). On the other hand, *Cf-Ecp4* was less prevalent by being present in only 6 accessions from 5 species (*S. lycopersicum*, *S. chmielewskii*, *S. arcanum*, *S. neorickii* and *S. pennellii*) (Figure 3). These are in addition to previously identified sources of *Cf-Ecp4* in *S. pimpinellifolium* (LA1683 and LA1689) (LAUGÉ *et al.* 2000). These results showed that both *Cf-Ecp5* and *Cf-Ecp4* are conserved within the wild tomato (*Solanum*) genus.

**FIGURE 3.**
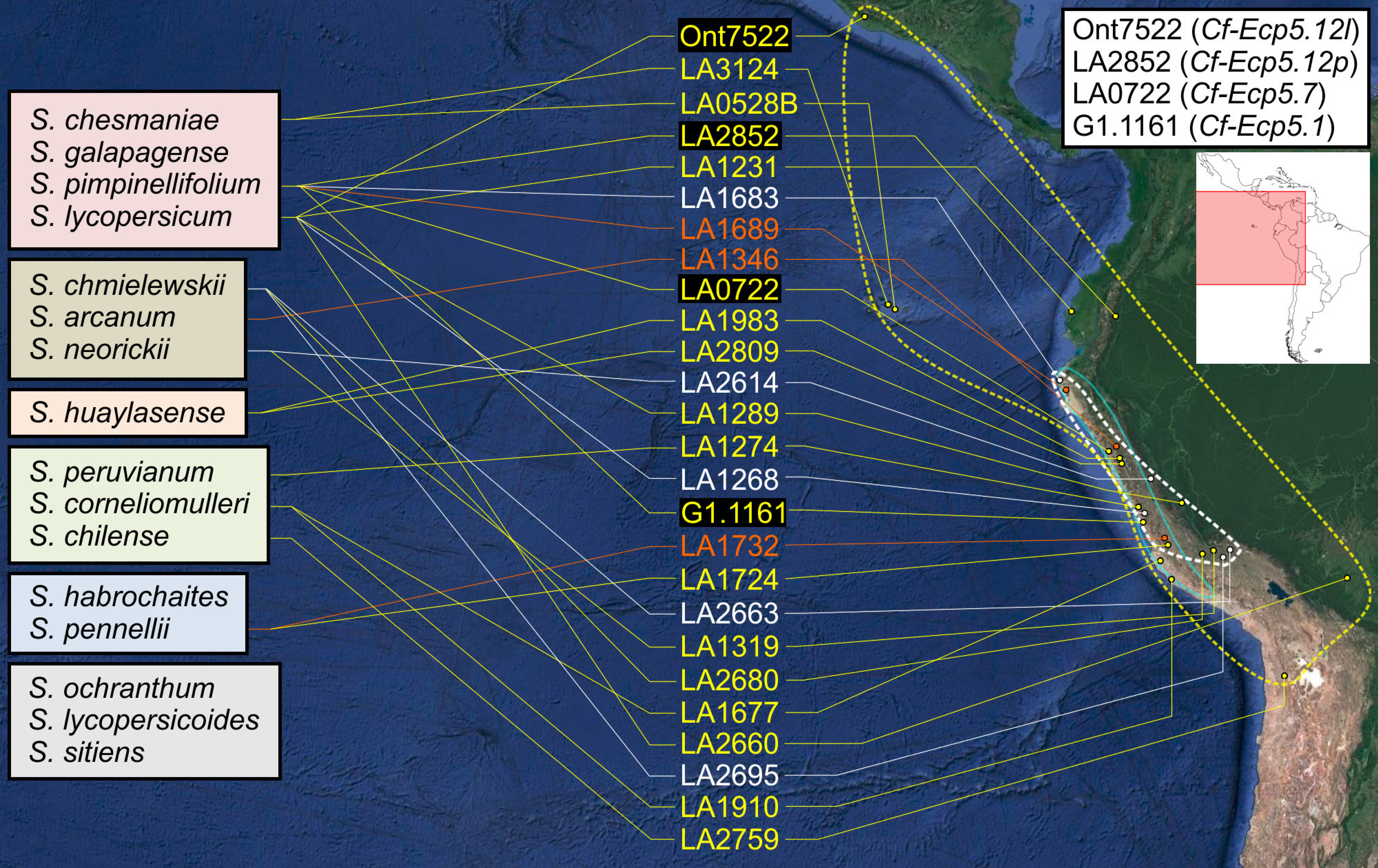
Geographic distribution of Ecp4- and Ecp5-induced HR on the north-western area of South America. The distribution of Ecp4- (white script) and Ecp5-responsive (yellow script) accessions following an agro-infiltration of seedlings with *PVX*:*Ecp4* and *PVX*:*Ecp5* respectively on a collection of 139 wild tomato accessions is shown. Accessions responding to both Ecp4 and Ecp5 are shown in orange script. The figure also includes *S. pimpinellifolium* accessions tested in a previous work (LAUGÉ *et al.* 2000), which were used for mapping in this study (black highlight). The tomato species used in this screen are also shown on the left, clustered in groups that reflect their approximate phylogenetic relationship as established previously (KRUIJT *et al.* 2005; RODRIGUEZ *et al.* 2009). The approximate distribution of Ecp4- and Ecp5-responsive accessions is shown as white and yellow dashed colored borders respectively, along with the distribution of *Cf-9* alleles (blue colored border) from a previous study (KRUIJT *et al.* 2005). The *Cf-Ecp5* genes that were mapped in this study are shown along with their accessions on the top right. The responsive accessions are ordered by geographic latitude and their location was estimated by using the TGRC and CGN database in conjunction with “Google Maps” online service (overview of map on the top right). Source material for breeding lines Ont7522 and G1.1161 were PI124161 and CGN15529 respectively, for which their collection sites are used in this map to locate the origin of their corresponding genes (KANWAR *et al.* 1980; LAUGÉ 1999; LAUGÉ *et al.* 2000).

### The tomato *Cf-11* and *Cf-3* genes are genetically linked on the top arm of chromosome 11

*S. lycopersicum* line Ont7716 carries the *Cf-11* gene, introgressed from *S. pimpinellifolium*, and its location has not been previously reported. Ont7716 is known to carry a copy of *Cf-4* as well (ENYA *et al.* 2009), so any mapping effort required an Ont7716 progeny line carrying *Cf-11* only to facilitate disease resistance phenotyping and subsequent mapping. To this end, 91 plants from a (Cf0 x Ont7716) x Cf0 population were inoculated with *C. fulvum* race 4 GUS, which can overcome resistance conferred by *Cf-4*, but not *Cf-11* (OLIVER *et al.* 1993). Results confirmed the presence of *Cf-11* (43:48 Resistant:susceptible, *p*-value(χ^2^) = 0.711) and resistant plants were randomly selected to be selfed. Seven selfed populations were infected with *C. fulvum* race 5 that cannot overcome resistance to either *Cf-4* or *Cf-11* (Table S4), thus any population segregating phenotypically in a 3:1 ratio would carry the *Cf-11* gene only. Four F_2_ families were identified that carried *Cf-11* only and further selfing and progeny tests (J476) resulted in the identification of a *Cf-11* homozygous plant, designated ‘Cf11N’ that was used as parent for mapping. However, preliminary attempts to identify molecular markers linked to *Cf-11* using Ont7716 x *S. pennellii* F_2_ population were unsuccessful due to challenges in disease phenotyping. Thus, to select a parent that will be polymorphic to Ont7716 for AFLP analysis and subsequent mapping of *Cf-11*, Ont7716 was analysed by DNA gel blots with probes that detect *Hcr9s* and *Hcr2s* (Figure S4A, S4B). From the results the Cf2 haplotype was chosen as a polymorphic parental line to be crossed with Ont7716. Resistant and susceptible F_2_ bulks from a (Cf2 x On7716) x Cf0 cross were analysed by AFLP analysis with 264 primer combinations and one marker (M1) linked *in cis* to *Cf-11* was identified. BLAST analysis on the M1 sequence showed that the marker is located at 1.232 Mb on chromosome 11 (based on the *S. pennellii* genome sequence) (File S1). To construct a genetic map, 938 progeny from J476 (Cf0 x Ont7716 background, segregating for *Cf-11*) were infected with *C. fulvum* race 4 GUS and 214 susceptible plants were genotyped with markers M1 and SSR136 (704:234 resistant:susceptible, *p*-value(χ^2^) = 0.979) (Table S5). The results positioned the locus between the two markers (M1 and SSR136), while a subsequent screen with markers 0124F20 and 136SP6 confirmed this and further delimited *Cf-11* to a 535 kb region (1.322-1.857 Mb) (Figure 4).

**FIGURE 4.**
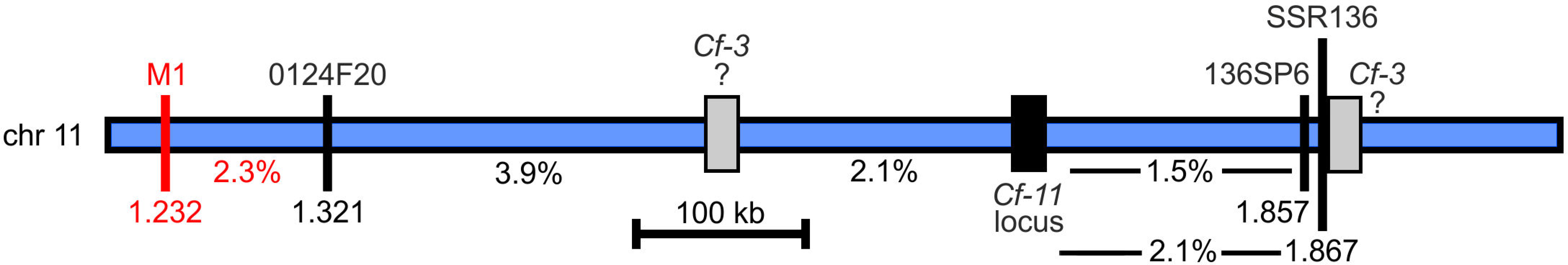
The *Cf-11* and *Cf3* loci are linked on the top arm of chromosome 11 in tomato. Genetic map of the chromosome 11 region at the *Cf-11* locus in Ont7716 (blue box). The interval was delimited by genotyping 201 susceptible plants from a population of 831 J476 progeny (Cf0 x Ont7716 BC_3_; following infection with C. *fulvum* race 4 GUS) with two markers, 0124F20 and 136SP6, and combining the data with previous results with another two markers (M1 and SSR136). The Cf-11 locus showed 3.9% recombination with marker 0124F20 (16/402 recombinant gametes) and 1.5% with 136SP6 (6/402 recombinant gametes). Distances are shown in megabases (Mb) and the location of each marker on the chromosome is also shown as Mb coordinates. The recombination fraction (%) between the *Cf-11* locus (black box) and each marker is shown, along with the hypothetical distal and proximal locations of the *Cf-3* locus (grey box) with 2.1% on either side of the Cf-11 locus.

The tomato genetic map indicated the *S. lycopersicum Cf-3* gene is also located on the top arm of chromosome 11 (KANWAR *et al.* 1980). Preliminary marker analysis showed that the Cf3 haplotype was similar to Cf11N at the *Cf-11* locus and therefore, these genes may be linked, or possibly allelic. The progeny of a (Cf3 x Cf11N) x Cf0 cross were inoculated with *C. fulvum* race 4 GUS that is restricted by either gene. If *Cf-3* and *Cf-11* are allelic, no susceptible progeny should have been recovered from this cross. However, a number of progeny (6 out of 288) were scored as susceptible, showing similar levels of fungal sporulation to Cf0 control plants at 14 dpi. Molecular analysis of these individuals confirmed they contained markers of the *Cf-3/Cf-11* haplotype, hence *Cf-3* and *Cf-11* appear to be closely linked (Figure 4). The chromosomal region where both *Cf-3* and *Cf-11* reside appears to be devoid of RLPs in the *Solanum lycopersicum* genome (JUPE *et al.* 2013; ANDOLFO *et al.* 2014; WEI *et al.* 2016; KANG AND YEOM 2018).

## DISCUSSION

### Expanding the *Cf* genetic map in tomato

Breeding for resistance to *C. fulvum* has a long history, but breeding for durable resistance will depend on successful introduction of novel *Cf* genes from wild tomato species (BAILEY 1947; KERR AND BAILEY 1964). The *Cf* gene map has been extended in this study to include new loci on three different chromosomes (Figure 5). The genetic mapping of *Cf-Ecp5.1* to *MW* further highlights the importance of this locus in the tomato genome to generate effector recognition specificities (JONES *et al.* 1994; THOMAS *et al.* 1997; TAKKEN *et al.* 1999; PANTER *et al.* 2002; YUAN *et al.* 2002; RIVAS AND THOMAS 2005; SOUMPOUROU *et al.* 2007; DE WIT *et al.* 2009; ZHAO *et al.* 2016). The reassignment of *Cf-Ecp5.1* at *MW* from *AU* is possibly due to the different functional analyses performed to assay *Cf-Ecp5* function. In this study we used only *PVX:Ecp5*, while in the original study a combination of *C. fulvum* infections and *PVX*:*Ecp5* were used in mapping that could have resulted in assaying two distinct, but linked fungal resistance genes (LAUGÉ *et al.* 2000).

**FIGURE 5.**
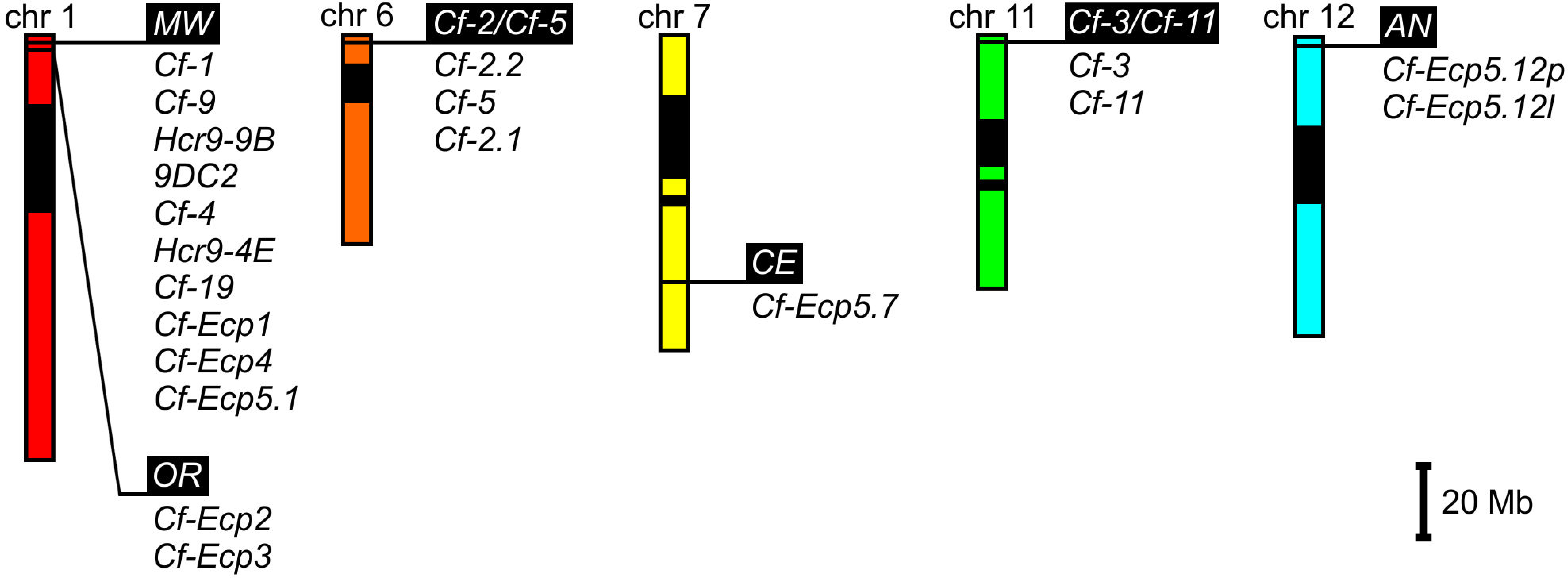
The expanded tomato *Cf* gene map. Schematic representation of tomato chromosomes 1, 6, 7, 11 and 12 as genetic maps that include all *Cf* genes mapped to date. Loci are shown relative to the physical map of each chromosome (based on the tomato physical map at Sol Genomics Network). Heterochromatic regions are shown as black boxes on each chromosome. Loci are shown as white script on black background with all their known *Cf* genes underneath. Scale is shown in megabases (Mb).

All previously characterized *Cf* genes on tomato chromosomes 1 and 6 encode RLPs (WULFF *et al.* 2001; WULFF *et al.* 2004). To accurately identify candidate RLP genes on the newly mapped tomato loci *CE, Cf-11/Cf-3* and *AN* on chromosomes 7, 11 and 12 respectively, a high quality genome sequence is needed from each accession, as *Cf* loci are highly polymorphic (WULFF *et al.* 2009). In an early attempt, we analysed the *S. lycopersicum* ‘Heinz 1706’ tomato genome sequence (CONSORTIUM 2012) and the *S. pimpinellifolium* LA1589 draft genome sequence (THE TOMATO GENOME *et al.* 2012) in regions delimited by the flanking markers for each locus in this study, retrieving some informative data only for ‘Heinz 1706’ (Table S7). However, recent studies have identified candidate *R* genes in the regions covering the novel *Cf* loci reported in this study (Table S7) (JUPE *et al.* 2013; ANDOLFO *et al.* 2014; WEI *et al.* 2016; KANG AND YEOM 2018). The *CE* and *Cf-11/Cf-3* loci, on chromosomes 7 and 11 respectively, reside in regions poor in RLPs (KANG AND YEOM 2018), while the *AN* locus resides in a hotspot for RLPs similar to the *Milky Way* locus on chromosome 1, which is confirmed by both our analysis (Table S7) and another study (KANG AND YEOM 2018). It will be interesting to know whether all these novel *Cf* genes encode RLPs or something else.

### The multiplicity of Ecp5 responses in tomato species

In some plant-microbe interactions, two or more loci may control the resistance phenotype in an extension of Flor’s gene-for-gene hypothesis (FLOR 1947; BOURRAS *et al.* 2016). However, these genes do not convergently evolve in different chromosomal locations, with the exception of the potato *Rpi-blb3* and *Rpi-mcq1* genes to date (AGUILERA-GALVEZ *et al.* 2018), which exhibit differential effector-specific resistance to Avr2-carrying *Phytophthora infestans* isolates (AGUILERA-GALVEZ *et al.* 2018). This distinctiveness of these two potato *R* genes recognizing Avr2 has been attributed to independent evolution of the underlying recognition mechanism, since the genes originate from Mexico (*Rpi-blb3*) and from Peru (*Rpi-mcq1*) (AGUILERA-GALVEZ *et al.* 2018). Avr2 is also a highly diverse effector within *Phytophthora* species (VLEESHOUWERS *et al.* 2011), which could facilitate convergent evolution of two *R* genes through overlapping recognition specificities.

In this study, from the mapping attempts on only four accessions, four *Cf-Ecp5* loci were mapped on three different chromosomes, exhibiting differences in HR strength in both *PVX*:*Ecp5* agro-infiltration assays and crosses to Ecp5-expressing transgenic tomatoes (Figure S3), while allelism crosses between different *Cf-Ecp5*-carrying accessions revealed putative suppression elements that point to distinct evolution of effector recognition or defense activation mechanisms in tomato (Table 2). Crosses between *S. pennellii* LA0716 and *S. pimpinellifolium* LA2852 or *S. lycopersicum* Ont7522 resulted in suppression of the Ecp5-induced HR at the F_1_ generation, while the same phenotype was also observed in a *S. pimpinellifolium* G1.1161 x (Cf0 x LA2852) background (Table 2). The most interesting example of Ecp5-induced HR suppression was observed in a cross between *S. lycopersicum* Ont7522 and F_1_ progeny from Cf0 x *S. pimpinellifolium* LA2852, in which both accessions carry a *Cf-Ecp5* gene on chromosome 12 at *AN* locus (Table 2). We genotyped 39 randomly selected wild-type plants to investigate if the Ecp5-induced HR suppression assorts independently of the *AN* locus on chromosome 12. The results indicated that the genetic elements responsible for this *Cf-Ecp5-*induced HR suppression were genetically unlinked to the *AN* locus and accession-specific for each *Cf-Ecp5* gene (Table S6). These data suggest that these *Cf-Ecp5* genes have evolved distinct HR-regulating elements.

Just like other characterized *Cf* genes, *Cf-Ecp5*s possibly also encode RLPs, which would lack any intracellular signalling domain and they may interact with the receptor-like kinase (RLK) SOBIR1 (LIEBRAND *et al.* 2013), or a similar RLK that will stabilize them, and require the RLK BAK1 for downstream immune signalling (VAN DER BURGH *et al.* 2019). However, functional tests will be required to determine if Ecp5 is perceived by functionally similar/distinct CfEcp5 proteins and whether they all require SOBIR1 and BAK1 for defense activation. The suppression phenotypes of Ecp5-dependent HR in different *Cf-Ecp5*-carrying accessions (Table 2) may also help to elucidate this question. Ecp5 is the most monomorphic effector reported in *C. fulvum*, having only one variant with a single polymorphic nucleotide in its single intron (STERGIOPOULOS *et al.* 2007; DE WIT *et al.* 2009), which raises an interesting question as to how these different accessions evolved genetically-distinct Ecp5 recognition specificities. All Ecp5-responding accessions originated from different geographic regions (Figure 5) (KANWAR *et al.* 1980; LAUGÉ 1999; LAUGÉ *et al.* 2000; RIVAS AND THOMAS 2005; DE WIT *et al.* 2009) and whether geographic origin plays a role in the evolution of strong responses or not remains to be seen. The geographic distribution of Ecp5-responding accessions appeared much wider than either *Cf-Ecp4* or *Cf-9* (Figure 3), which could be explained by the convergent evolution of multiple distinct *Cf-Ecp5* loci under different environmental conditions or low selection pressure (STERGIOPOULOS *et al.* 2007; BOLTON *et al.* 2008). Another interesting question is how many distinct *Cf-Ecp5* loci and types of Ecp5-responses exist throughout tomato species that were not screened in our study (Figure 3) (Table S3). If *Cf-Ecp5* genes are to be incorporated into breeding programs, their allelic and variant interactions need to be studied further in different backgrounds and with multiple *C. fulvum* strains.

### Do effector-specific *R* genes emerge and evolve anywhere?

*Cf* genes have been repeatedly bred into cultivated tomato from wild relatives such as *S. pimpinellifolium*, *S. hirsutum* and *S. peruvianum* (G. ATHERTON AND RUDICH 1986) and according to a recent study (KANG AND YEOM 2018) most of these types of genes (RLPs) are located on tomato chromosomes 1 and 12. Some regions appear to be the source of major *Cf* loci, like *MW*, which are able to generate multiple effector-specific RLPs (RIVAS AND THOMAS 2005; DE WIT *et al.* 2009). Although distinct recognition of multiple effectors by one or more R proteins is known (MACKEY *et al.* 2002; MA *et al.* 2018), the opposite has only been observed as convergent evolution in different species (ASHFIELD *et al.* 2014; SAUCET *et al.* 2015; CARTER *et al.* 2018) with only one recent example in closely related species (AGUILERA-GALVEZ *et al.* 2018). However, considering the monomorphic nature of Ecp5 and level of genetic complexity observed amongst *Cf-Ecp5* genes in this study, we have to address this slight paradox in terms of evolution. There are hypotheses that might be able to explain the *Cf-Ecp5* evolution, since conventional conservation of a *Cf-Ecp5* allele through natural selection (MAURICIO *et al.* 2003) is unlikely to be the source of four genetically distinct genes, nor are the putative overlapping specificities of distinct R proteins for a highly variable effector like Avr2 (GILROY *et al.* 2011; AGUILERA-GALVEZ *et al.* 2018), as Ecp5 is not.

The first hypothesis involves small-scale genomic duplications following unequal crossover events or other chromosomal anomalies that have allowed the dispersal of an original cluster harbouring the ancestral *Cf-Ecp5* gene throughout the tomato genome prior to speciation (BAUMGARTEN *et al.* 2003; MEYERS *et al.* 2005; RENSING *et al.* 2008; FLAGEL AND WENDEL 2009). The second hypothesis involves transposons and other reverse-transcriptase-mediated duplication that can lead to ‘macro-transposition’ of *Cf-Ecp5* genes or their clusters (FREELING *et al.* 2008; HUANG AND DOONER 2008; XIAO *et al.* 2008). The third hypothesis for the *Cf-Ecp5* evolution could be sought in the mechanisms involved in plant *R* gene creation, which is the moment pseudogenes or paralogues become fully functioning resistance genes capable of activating defenses. A critical factor in addressing this is probably the long-term effects of biotic stress on DNA evolution and *R* gene loci structure, which have only been investigated partially and indirectly (KOVALCHUK *et al.* 2003; BIÉMONT AND VIEIRA 2006; BOYKO *et al.* 2007; BOYKO *et al.* 2010). Nevertheless, sequencing and bioinformatic analysis of all *Cf-Ecp5* genes and their flanking sequences will uncover the nature of these *R* genes and help to understand their function.

### Conclusions

In plant pathology, it is important to consider the underlying complexity of each pathosystem and its gene-for-gene interactions. The discovery of novel *Cf* loci in this study and expansion of the *Cf* repertoire with loci of variable HR strength (*Cf-Ecp5s*) has potential implications for breeding plant disease resistance. Vertical resistance in tomato can be defeated quickly as strains of *C. fulvum* that overcame several *Cf* genes have been reported already (KERR AND BAILEY 1964; LUDERER *et al.* 2002; ENYA *et al.* 2009). *C. fulvum* secretes approximately 70 apoplastic proteins *in planta* (MESARICH *et al.* 2017), from which a significant number act as effectors and are recognised in wild tomato accessions, making it very likely that a large number of novel *Cf* genes can be exploited for breeding purposes in the future. Considering that Ecp5 is a core effector in *C. fulvum* and has no variability, studying the differences between distinct *Cf-Ecp5* genes in the tomato germplasm can facilitate our understanding on effector- or strain-specific recognition and defense activation, while allowing us to choose or engineer optimal *Cf* gene variants that have the strongest possible defense responses to different *C. fulvum* strains with minimal cost-to-fitness. Furthermore, the simultaneous stacking of different variants of *Cf-Ecp5* genes into one cultivar could extend the duration of resistance to *C. fulvum* and constitute a model for horizontal resistance in other plant pathosystems in the future (DANGL *et al.* 2013).

## ACKNOWLEDGEMENTS

We thank James Brown, Jonathan Jones, Brande Wulff, Mark Coleman, and Farid El Kasmi for advice and suggestions. Both M.I. and E.S. were supported by the Greek State Scholarships Foundation (IKY) awards for postgraduate study. Research with recombinant *PVX* was authorized under DEFRA licence PHL 170/5467.

